# The Hurst exponent as a marker of inhibition in the developing brain

**DOI:** 10.1101/2024.09.29.615675

**Authors:** Monami Nishio, Monica E. Ellwood-Lowe, Mackenzie Woodburn, Cassidy L. McDermott, Anne T. Park, Ursula A. Tooley, Austin L. Boroshok, Joanes Grandjean, Allyson P. Mackey

**Author notes:** Correspondence (A.P.M), Address: Department of Psychology, School of Arts and Sciences, University of Pennsylvania, Philadelphia, Pennsylvania 19104, United States.

## Abstract

The maturation of inhibitory neurons is crucial for regulating plasticity in developing brains. Previous work using computational models has suggested that the Hurst exponent, the decay in power over frequency, reflects inhibition, but empirical data supporting this link is sparse. Here, we took a cross-species approach to validating the Hurst exponent of fMRI as a marker of inhibition, then characterized the development of the Hurst exponent in childhood. We found significant spatial correlations between the Hurst exponent and ex vivo parvalbumin mRNA expression in human children and adults, and between the Hurst exponent and parvalbumin-positive cell counts in mice. We identified a plateau in the mRNA expression by late childhood, aligning with the Hurst exponent plateau in both humans and rats. In sum, this work suggests that the Hurst exponent can be used to study the development of inhibition *in vivo*, and in the future, to understand individual differences in plasticity.

## Introduction

Children’s brains are incredibly plastic, allowing them to rapidly learn and refine new skills. Converging evidence from human and animal neuroscience points to the importance of inhibitory cells for regulating plasticity^1^. Inhibitory cells adjust the timing and strength of synaptic inputs, which is crucial for triggering spike-timing-dependent plasticity^1^. However, inhibition in the brain has been hard to measure in vivo. As a result, the precise timing of changes in inhibition and its variation across cortex is unknown. Elucidating the process of inhibitory development is critical for understanding healthy brain development and identifying windows of developmental vulnerability and opportunity.

Inhibition is orchestrated by GABAergic (Gamma-aminobutyric acid) cells in cortex^2^. Most inhibitory neurons can be classified according to their expression of three proteins: parvalbumin (PV+, 40% of GABAergic cells), somatostatin (SST+, 30%), and vasoactive intestinal polypeptide (VIP+, 10-15%)^3^. These three classes have distinct synaptic and functional features that differentially modulate cortical activity^4^. PV+ cells form synapses on perisomatic regions of excitatory pyramidal cells, effectively inhibiting their spiking, resulting in shorter activation timescales to facilitate the processing of quickly changing information such as sensorimotor inputs^5,6^. PV+ cells are found in higher numbers in sensory and motor cortical regions than in association cortex^5,7^. SST+ cells target dendrites of cortical projection neurons^4^ to help filter out distracting or irrelevant input signals^5,8^, and are found in higher numbers in association cortex than in sensory and motor regions^5,7^. VIP+ cells inhibit SST+ and PV+ cells, serving as a disinhibitory mechanism^9^, and are evenly distributed across cortex^5^.

Manipulations of PV+ cells lead to alterations in developmental plasticity^10–13^. For example, accelerating the maturation of PV+ cells in visual cortex by removing polysialic acid leads to an earlier onset and closure of the critical period for ocular dominance^11^. In contrast, destroying peri-neuronal nets around PV+ cells reactivates visual cortical plasticity in adult animals^14^. SST+ and VIP+ cells also play a role in regulating plasticity, primarily in adulthood. For instance, ontogenetically activating VIP+ cells to suppress SST+ cells enhances plasticity in the brains of adult mice^15^.

Postmortem studies in several species have characterized the protracted maturation of PV+ cells. For instance, in rats, PV levels plateau around adolescence (postnatal day [PD] 40) in prefrontal cortex^16,17^. In macaques, the adult laminar pattern of PV+ cell distribution is reached between three and five months of age in the inferior temporal, posterior parietal and prefrontal cortices^18^. In humans, the expression of PV mRNA plateaus around the age of five to ten years old in dorsolateral prefrontal cortex^19,20^. However, it is difficult to build a model of how inhibition matures based on mRNA expression because pediatric postmortem human samples are thankfully rare, and these samples often show clinical abnormalities. Moreover, these studies are typically focused on only a handful of brain areas, making it challenging to track developmental trajectories across cortex.

To fully understand how inhibition develops in humans, we need developmentally appropriate in vivo non-invasive measures. One approach is magnetic resonance spectroscopy (MRS)^21^. MRS can be used to obtain measures of the concentration of GABA. However, age-related changes in GABA showed inconsistent results across different sub-regions of prefrontal cortex and across hemispheres^22^. Furthermore, MRS can only sample one large voxel at a time and is sensitive to motion artifacts, limiting its use for whole-brain analysis and developmental studies^23,24^. Additionally, MRS data are not widely available in large public datasets, further complicating efforts to study developmental changes in inhibition comprehensively.

One strong contender for measuring inhibition in vivo in humans is the Hurst exponent. For a spectral density of the form 1/f α, the Hurst exponent α measures the slope of the log-transformed spectral density and hence characterizes the power-law scaling of the amplitude fluctuations in various types of neuroimaging time series, including EEG and fMRI BOLD signals^25^. 2α + 1 is equal to the aperiodic exponent, which has also recently been introduced as a measure of plasticity^26^. In computational models, increasing inhibition leads to an increase in the Hurst exponent of simulated local field potential and blood oxygen level dependent (BOLD) signals^27–29^. In mice, chemogenetically reducing the excitability of pyramidal neurons leads to an increase of the Hurst exponent of the recorded LFP signal^27^. In adolescent humans, the Hurst exponent derived from electroencephalogram (EEG) data correlates with a MRS measure of glutamate/GABA balance in dorsolateral prefrontal cortex^30^. A few studies have characterized developmental changes in the Hurst exponent. In mice, the Hurst exponent of the local field potential increases during the initial two weeks of life^27^. Similarly, in humans, the Hurst exponent of the EEG signal increases in the first few weeks of life^27^. However, to date, no studies have characterized changes in the Hurst exponent in childhood.

If the Hurst exponent of the fMRI signal is indeed a valid measure of inhibition, and therefore plasticity, it could be used to characterize how plasticity varies across cortical areas in the developing brain. Recent studies on the human brain have unveiled a hierarchical development of various structural and functional aspects along a sensorimotor-association (S-A) axis^31^. The timing of critical periods has been proposed to vary along the S-A axis, with sensory areas undergoing early critical periods and association areas showing critical periods in adolescence^32^. Consistent with this hypothesis, the distribution of PV+ cells fully matures in macaques first in primary visual cortex, followed by secondary visual cortex, and then by higher-order regions of the inferior temporal, posterior parietal, and prefrontal cortices^18^. However, there are currently no studies that have demonstrated a hierarchy of inhibitory development in humans.

In the present study, we employed a cross-species, multi-method approach to characterize the development of inhibition in childhood. First, we investigated the spatial correlation between the Hurst exponent, calculated from resting-state fMRI data, and inhibitory marker mRNA expression in human children and adults. Next, we replicated this correlation in fMRI and cell density data from mice. We then traced the developmental trajectories of the Hurst exponent in human children, mice, and rats. Together, this work supports the use of the Hurst exponent to study the development of inhibition across cortex.

## Results

### The Hurst exponent correlates with parvalbumin mRNA expression across cortex

We calculated the Hurst exponent of resting state fMRI data in children (ages 4-11 years, n = 128) and adults (ages 18-25 years, n = 46). We parcellated cortex into 400 equally-sized regions^33^. Using the Fourier transform, we computed the power spectral density (PSD) of the time series of each cortical parcel and estimated the slope of the best-fit line to the PSD. A higher Hurst exponent, or a steeper slope, is thought to reflect more inhibition because there is less power at high frequencies (Figure 1A). The topography of the Hurst exponent across cortex in children and adults, and its distribution along the sensorimotor-association axis, is shown in Figure S1.

**Figure 1.**
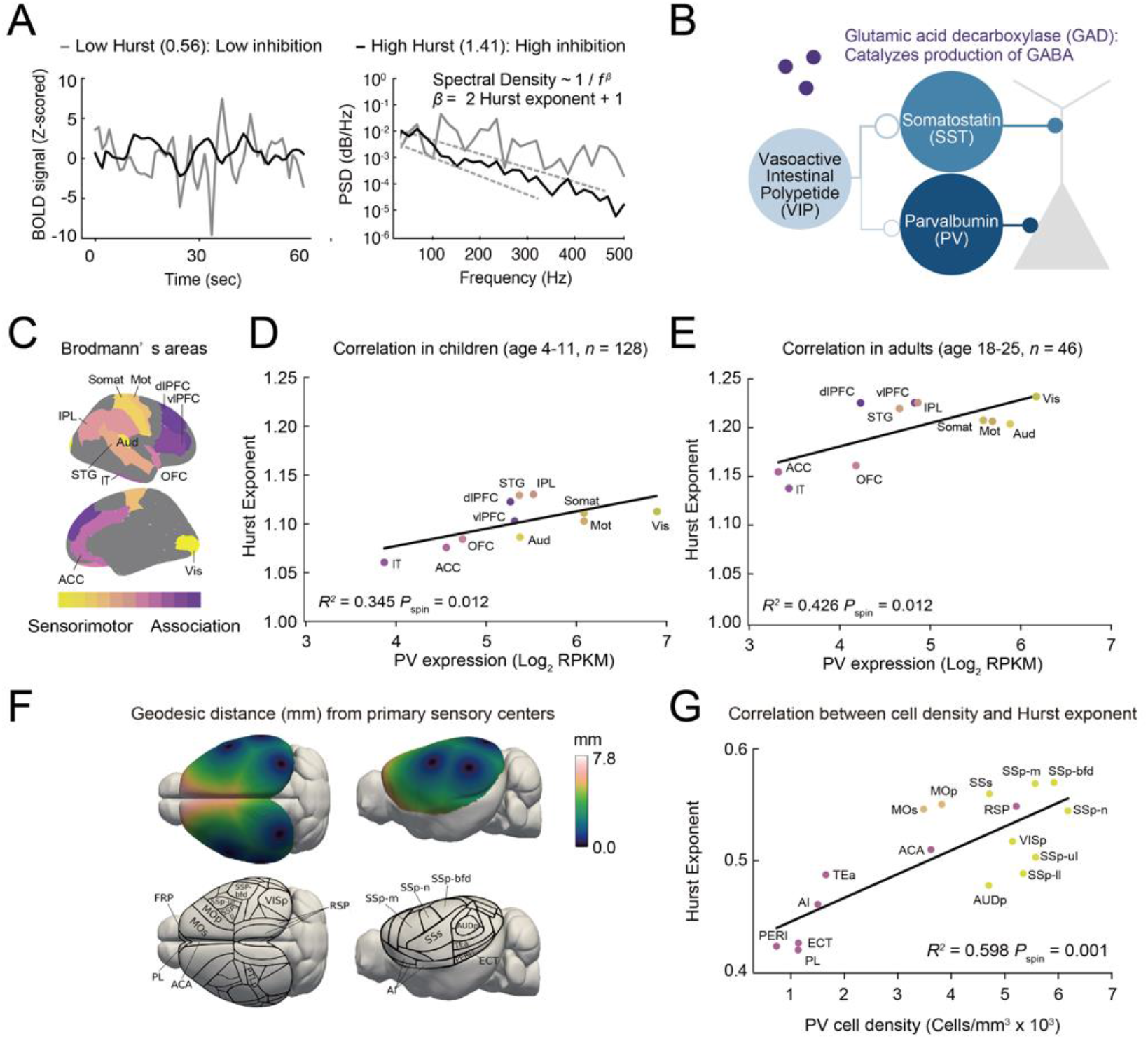
The Hurst exponent correlates with PV mRNA expression across cortex. **(A)** Examples of fMRI time series with high and low Hurst exponents and power spectrum densities (PSD). **(B)** Illustration of inhibitory neuron types, including parvalbumin (PV), somatostatin (SST), vasoactive intestinal polypeptide (VIP), as well as proteins involved in GABA production, glutamic acid decarboxylase (GAD 1 and 2). **(C)** Brodmann’s areas from which inhibitory marker expression was analyzed. Brain regions are arranged along a spectrum from sensorimotor (yellow) to association (purple).Vis; Visual Cortex (BA17), Aud; Auditory Cortex (BA41), Somat; Somatosensory Cortex (BA1-3), Mot; Motor Cortex (BA4), STG; Superior Temporal Gyrus (BA22), IPL; Inferior Parietal Lobe (BA39,40), OFC; Orbitofrontal Cortex (BA11-14), ACC; Anterior Cingulate Cortex (BA24,25), IT; Inferior Temporal Gyrus (BA20), vlPFC; Ventrolateral Prefrontal Cortex (BA44,45), dlPFC; Dorsolateral Prefrontal Cortex (BA8,9,46). **(D, E)** Correlation between PV expression (Log_2_ Reads Per Kilobase per Million mapped reads [RPKM]) and the Hurst exponent in children (D) and adults (E) (*R*^*2*^ = 0.345, *F(1, 9) = 6*.*272, P*_spin_ = 0.012 for children; *R*^*2*^ = 0.426, *F(1, 9)* = 8.415, *P*_spin_ = 0.012 for adults.). PV expression was extracted from Brodmann’s areas, and Hurst exponents were extracted from the Schaefer-400 parcels with more than 50% overlap with each Brodmann’s area. **(F)** Geodesic distance (mm) from primary sensory centers, which were defined as the geometric centers of the primary somatosensory, visual, and auditory areas (replicated from Huntenburg et al., 2021). Brain regions are arranged along a spectrum from closer (blue) to farther (pink) from the primary sensory centers. Anatomical segmentations are based on the Allen Mouse Brain atlas^36^. **(G)** Correlation between PV+ cell density and Hurst exponent. Sensory regions (yellow) include VISp; Primary visual area, AUDp; Primary auditory area, SSp-n; Primary somatosensory area, nose domain, SSp-m; Primary somatosensory area, mouth domain, SSp-bfd; Primary somatosensory area, barrel field, and SSs; Supplementary somatosensory area. Motor regions (orange) include MOp; Primary somatomotor area and MOs; Secondary somatomotor area. Assocation regions (purple) include TEa; Temporal association areas, ECT; Ectorhinal area, PERI; Perirhinal area, AI; Agranular insular area, RSP; Retrosplenial area, ACA; Anterior cingulate area, and PL; Prelimbic area. Linear regression analysis indicates a significant correlation (*R*^*2*^ = 0.598, *F(1, 15) = 24*.*840, P*_spin_ = 0.001).

To assess the validity of the Hurst exponent as an inhibitory marker, we investigated the correlation between the Hurst exponent and mRNA expression of markers of inhibitory neurons from the Brainspan Atlas of the Developing Human Brain (Figure 1B). We analyzed three markers of specific inhibitory neuron subtypes: parvalbumin (PV), somatostatin (SST), and vasoactive intestinal polypeptide (VIP). We also analyzed the expression level of glutamic acid decarboxylases (GAD) 1 and 2, which catalyze the production of GABA and are expressed across all types of inhibitory neurons. We included mRNA data from all 11 Brodmann’s areas^34^ sampled at ages 3, 8, 11, and 13 for children (n = 6) and ages 18, 19, 21 and 23 for adults (n = 4) (Figure 1C). As there were only two regions sampled for a 4-year-old, we excluded these data. No data were available for other ages. We aligned the Brodmann’s areas with the Schaefer 400 parcellation to extract Hurst exponents and to assign the regions to a sensorimotor-association axis rank. The significance of the correlation was assessed using spatially constrained rotation tests, known as ‘spin tests’^35^, because cortical data often exhibit distance-dependent spatial autocorrelation potentially inflating the significance of correlations between two cortical feature maps.

The distribution of the Hurst exponent significantly correlated with the expression of PV across cortex both in children (Figure 1D, R^2^ = 0.345, F(1, 9) = 6.272, P_spin_ = 0.012) and in adults (Figure 1E, R^2^ = 0.426, F(1, 9) = 8.415, P_spin_ = 0.012). Sensory and motor regions tended to have higher PV expression, as previously established^7^, as well as higher Hurst exponents. In contrast, the Hurst exponent showed a negative correlation with SST expression in adults but not in children. It also did not correlate with VIP expression, GAD1, or GAD2 for either children or adults (Figure S2).

Given that PV expression is notably higher in sensory regions compared to association regions, one possibility is that the correlation between the Hurst exponent and PV expression stems solely from their alignment along the S-A axis. If this is the case, we would expect similarly strong correlations between other genes aligned with the S-A axis and the Hurst exponent. We found 655 genes significantly correlated with the S-A axis in both children and adults (P < 0.05, uncorrected). However, of these 655 genes, only four were significantly correlated with the Hurst exponent (PVALB, KCNS1, JDP2, ITIH5). Among these, PV had a numerically higher R^2^ for both children and adults’ Hurst exponent compared to the other three genes. We also performed a partial correlation test and found that even when controlling for S-A axis rank, PV expression and the Hurst exponent still showed a significant correlation (children: r(9) = 0.730, P = 0.016; adults: r(9) = 0.811, P = 0.004). This suggests that their correlation cannot be solely attributed to their alignment with the S-A hierarchy. Another possibility is that although PV significantly correlates with the Hurst exponent, other genes might correlate even more strongly and better explain the spatial distribution of the Hurst exponent. Out of the 21,377 unique genes in our dataset, only 42 genes (< 0.2%) displayed significant correlations (P < 0.05 uncorrected) with both child and adult Hurst exponents (Figure S3). Among these 42 genes, only eight other genes (FNDC4, PIM3, F10, SNX21, UBE2QL1, SEZ6L2, RXRB, IL32) had a numerically higher R^2^ for both children and adults than PV. These eight genes encode proteins known to play key roles in metabolism and have no known roles in brain function. Therefore, we conclude that PV is still the most likely candidate to explain the spatial distribution of the Hurst exponent.

Next, we assessed whether the correlation between the Hurst exponent and PV+ cells extended to mice. In lieu of mRNA expression data, we used cell density (cells per mm3) information for PV+, SST+ and VIP+ cells^5^. The Allen Mouse Brain atlas^36^ was used for anatomical segmentation of the whole-brain cell type distribution. For the Hurst exponent calculation, we used a publicly available dataset of mouse resting-state fMRI (doi:10.18112/openneuro.ds001653.v1.0.2). We arranged brain regions of the Allen Mouse Brain atlas^36^ based on the geodesic distance from primary sensory centers, which were defined as the geometric centers of the primary somatosensory, visual, and auditory areas (Figure 1F, reproduced from Huntenburg et al., 2021^37^). Geodesic distance measures the shortest path between two points on a surface, following the surface’s contours^37^. Distances were calculated using the surface mesh representation of the Allen Mouse CCF v3^38^.

As in humans, we found relatively higher PV+ cell density and higher Hurst exponents in sensory and motor regions compared to association regions (Figure 1G). The Hurst exponent was strongly positively correlated with the density of PV+ cells (Figure 1G, R^2^ = 0.598, F(1, 15) =24.840, P_spin_ = 0.001). The Hurst exponent was negatively correlated with SST+ and did not correlate with VIP+ (Figure S4A and S4B). We did find that the anesthesia protocol significantly influences the correlations between the Hurst exponent and cell counts. We observed a significant positive correlation between PV+ cell density and the Hurst exponent in mice anesthetized with 1% isoflurane, but this correlation was eliminated under the Med/Iso (Medetomidine 0.05 mg/kg bolus/0.1 mg/kg/h infusion (i.v.) + Isoflurane 0.5%) anesthesia protocol (Figure S4C).

### Developmental plateaus in inhibition in humans, mice, and rats

We first characterized associations between mRNA markers of inhibition and age during childhood in humans across 11 cortical regions (Figure 2A). We observed increases in mRNA expression which plateaued by the age of eight for all inhibitory markers (Figure 2B-F, all ages greater than age 3, ANOVA P < 0.001, df_between_ = 3, df_within_ = 55). However, because no samples were available for children ages 4 to 7, the timing of the plateau is difficult to estimate. Associations with age did not differ across regions of cortex (P’s > 0.05).

**Figure 2.**
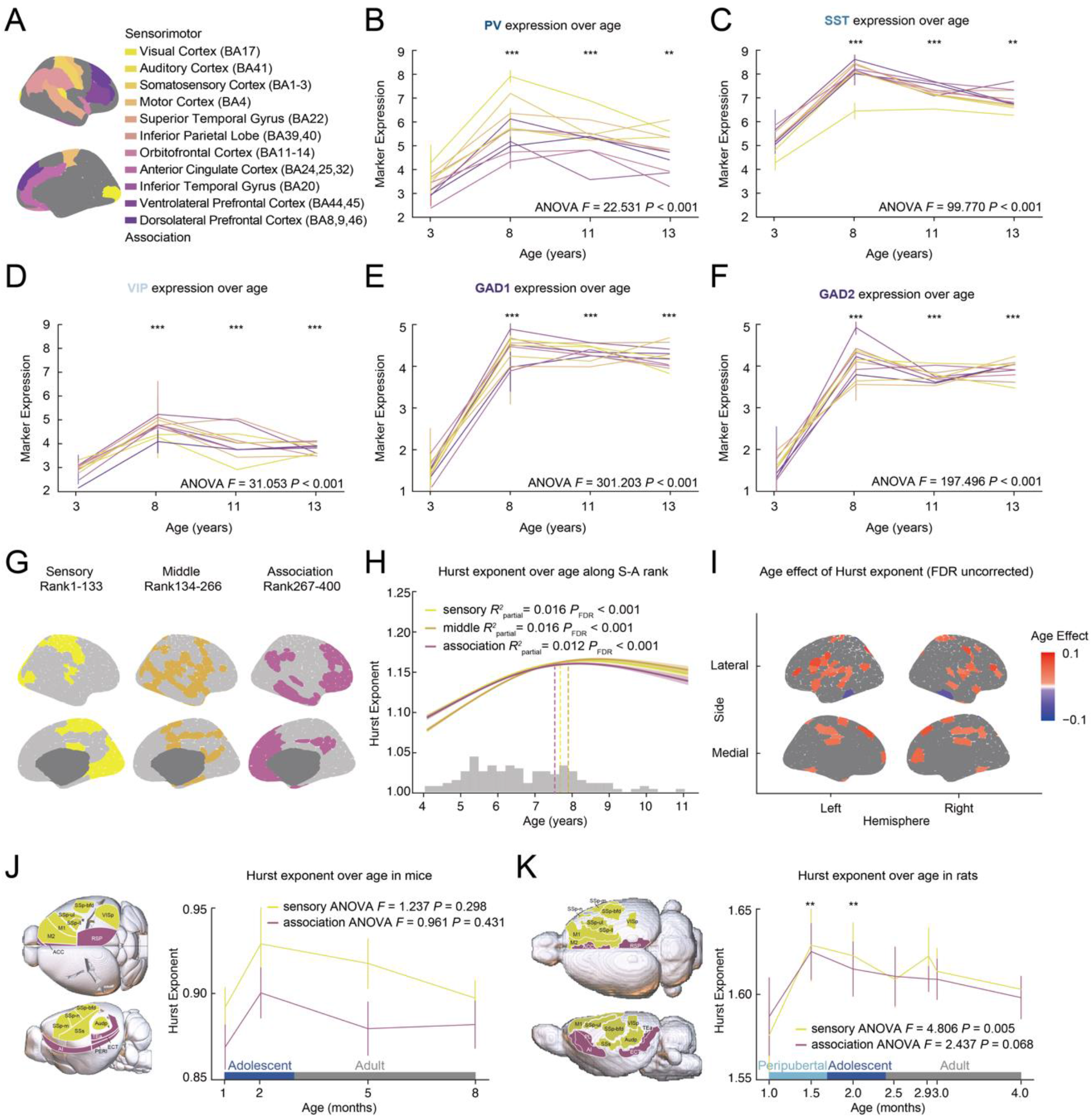
Plateau of inhibitory development in middle childhood. **(A)** Brodmann’s areas (BA) from which inhibitory marker expression was analyzed. Brain regions are arranged along a spectrum from sensorimotor (yellow) to association (purple). **(B-F)** Expression of PV (B, ANOVA *F(3, 55)* = 22.531, *P* < 0.001), SST (C, ANOVA *F(3, 55)* = 99.770, *P* < 0.001), VIP (D, ANOVA *F(3, 55)* = 31.053, *P* < 0.001), GAD1 (E, ANOVA *F(3, 55)* = 301.203, *P* < 0.001), and GAD2 (F, ANOVA *F(3, 55)* = 197.496, *P* < 0.001) across development from age 3 to 13. Line colors correspond to colors in (B). *** *P* < 0.001, posthoc Tukey HSD test comparing each older age group with 3-year-olds. Error bars represent ± *SD* across donors. **(G)** The 400 parcels are segmented along the Sensory-Association^31^ axis into three bins; sensory (yellow), middle (orange) and association (purple). Each bin contains 133 parcels. **(H)** GAM-predicted developmental trajectories of the Hurst exponent. Regional trajectories for each bin illustrate the GAM-predicted Hurst exponent values at each age, accompanied by a 95% confidence interval (sensory; partial *R*^*2*^ = 0.016, *P*_FDR_ < 0.001, middle; partial *R*^*2*^ = 0.016, *P*_FDR_ < 0.001, association; partial *R*^*2*^ = 0.012, *P*_FDR_ < 0.001). A dashed line marks the age at which the first derivative of the age smooth function (Δ Hurst exponent / Δ age) becomes insignificant. The age distribution of 128 participants (*M* = 6.55, *SD* = 1.36) is depicted as a histogram along the x-axis. **(I)** Associations between age and the Hurst exponent at *p* < .05 uncorrected. **(J)** The Hurst exponent across development of mice (sensory; ANOVA *F(3, 107)* = 1.237, *P* = 0.298, association; ANOVA *F(3, 107)* = 0.961, *P* = 0.431). Sensorimotor regions are shown in yellow, while association regions are shown in purple. **(K)** The Hurst exponent across development of rats (sensory; ANOVA *F* = 4.806, *P* = 0.005, association; ANOVA *F(6, 138)* = 2.437, *P* = 0.068). ** *P* < 0.01, posthoc Tukey HSD test comparing each older age group with one month-old. Error bars represent ± *SD* across subjects. Sensorimotor regions are shown in yellow, while association regions are shown in purple.

Next, we investigated whether a plateau in inhibition was also detectable in the Hurst exponent in children. We fitted region-specific generalized additive models (GAMs) with a smooth term for age. Each GAM estimated a smooth function (the model age fit) that described the relationship between the Hurst exponent and age, thereby modeling a region’s developmental trajectory. Gender, mean framewise displacement and the number of censored volumes were included as covariates; slopes did not differ by gender. We calculated the partial R^2^ change between the full GAM model and the reduced model (no age effects) and signed the partial R^2^ by the sign of the average first derivative of the smooth age function (effect direction). The 400 parcels were divided along the S-A axis into three bins, each containing 133 parcels (Figure 2G). Across sensory, middle, and association regions, the Hurst exponent plateaued around ages seven to eight years old (Figure 2H, sensory; partial R^2^ = 0.016, P_FDR_ < 0.001, middle; partial R^2^ = 0.016, P_FDR_ < 0.001, association; partial R^2^ = 0.012, P_FDR_ < 0.001; a dashed line indicates the age at which the first derivative of the age-smooth function [Δ Hurst exponent / Δ age] becomes insignificant). Age effects did not differ significantly across parcels by S-A rank. Areas where the Hurst exponent increased significantly across development are shown in Figure 2I.

To understand the developmental trajectory of the Hurst exponent in mice, we used the fMRI data obtained at one, two, five and eight months old, roughly corresponding to adolescence and young adulthood^39^. We divided cortex into sensorimotor regions (eight sensory regions, two motor regions) and association regions (seven regions). In aggregate, sensorimotor and association regions did not show significant age effects in this age range (Figure 2J, sensorimotor; ANOVA F(3, 107) = 1.237, P = 0.298, association; ANOVA F(3, 107) = 0.961, P = 0.431).

Since the developmental stage most similar to human childhood occurs before one month in mice, and mice younger than one month are difficult to image with fMRI, we conducted an additional developmental analysis using rat fMRI. We used a public rat fMRI dataset with data available for 1.0, 1.5, 2.0, 2.5, 2.9, 3.0 and 4.0 months ^40^, an age range that roughly corresponds to peripubertal (1.0, 1.5 months), adolescence (2.0 months), and adulthood (2.5-4.0 months)^41^. The SIGMA anatomical atlas^42^ was employed for whole-brain parcellation. Within this age range, sensory regions showed a significant age effect, while association regions exhibited a trending age effect (Figure 2K, sensorimotor; ANOVA F(6, 138) = 4.806, P = 0.005, association; ANOVA F(6, 138) = 2.437, P = 0.068). The Hurst exponent at 1.5 and 2.0 months was significantly higher than at 1.0 month according to the post hoc Tukey HSD test in sensorimotor regions (1.5 month; q(18, 20) = −0.032, P = 0.019, 2.0 month; q(76, 78) = −0.028, P = 0.003). From 1.0 to 1.5 month, there was a significant increase of the Hurst exponent for both sensorimotor and association regions (sensorimotor; t(18) = −2.742, P = 0.013, association; t(18) = −2.124, P = 0.048, independent t-test).

### Sensitivity analysis

We conducted our main analyses with more stringent motion exclusion criteria (mean framewise displacement < 0.5mm) due to the significant correlation we observed between the motion (mean framewise displacement and number of censored volumes) and the Hurst exponent in children (Figures S5A, B). No adults were excluded under this more stringent criterion, but 22 children were dropped from analyses. The relationship between PV expression and the Hurst exponent was significant in the smaller sample of children (R^2^ = 0.281, F(1, 9) = 4.910, P_spin_ = 0.025). Developmental trajectories followed a similar shape (Ps < .001 for sensory, middle, and association regions) but the timing of the plateau appeared slightly later (Figure S5C).

## Discussion

We employed a cross-species, multi-modal approach to investigate the early development of inhibitory circuits. We found significant spatial correlations between the Hurst exponent and the expression of parvalbumin (PV) across cortex in humans and in mice. We also found that PV expression and the Hurst exponent show plateaus by late childhood across sensorimotor and association areas. These results support the use of the Hurst exponent as a non-invasive method to investigate the development of inhibition, and by extension, plasticity. In this study, we discovered a correlation between the Hurst exponent and the post-mortem expression of PV mRNA in human cortex. We also found that the Hurst exponent correlates with the density of PV+ cells across cortex in mice. In both species, the Hurst exponent was more correlated with PV cells than other inhibitory subtypes such as SST+ and VIP+ cells. This variation between inhibitory subtypes might be explained by the different functions of each subtype^4^. PV+ cells predominantly form synapses on perisomatic regions, exerting significant influence over the output of pyramidal neurons^4^. In contrast, SST+ cells primarily target dendrites to regulate inputs, while VIP+ cells synapse with both PV+ and SST+ cells, thereby not directly regulating either the input or output of pyramidal cells^8,43^. Given that PV+ cells are the critical regulators of plasticity in developing brain^10–13,44^, this finding highlights the potential of the Hurst exponent as a promising in vivo marker for measuring children’s brain plasticity.

We observed a plateau in inhibition by the age of eight years in both the mRNA and fMRI human data. This result is consistent with a previous study examining post-mortem human samples of dorsolateral prefrontal cortex, which noted an increase in PV gene expression from neonates to middle childhood, followed by a slight decrease in late childhood^20^. In human primary visual cortex, another metric of inhibition, the balance of GABAAα1: GABAAα2 receptor subunits reaches adult level at the age of four to five years old^45^. The expression level of gephyrin, a GABAA receptor anchoring protein, also shows a peak in expression during childhood (5-11 years old)^45^. This result is also consistent with a wide variety of behavioral studies showing that skills such as reasoning, executive function, emotion regulation, and social cognition show significant improvement until around eight to ten years old, after which the rate of change decreases^46–49^. Our finding that inhibition plateaus around middle childhood suggests that this stabilization of inhibition may contribute to the slowdown in skill acquisition.

Contrary to what we initially expected, we did not find a clear hierarchical relationship along the S-A axis in the development of both mRNA expression and the Hurst exponent. Recent studies have shown various events in cortical morphogenesis follow the S-A hierarchy^31,32^. However, certain critical events, such as astrogliogenesis^50^ or the rapid proliferation phase of cortical synaptogenesis^50–54^, occur simultaneously throughout cortex. In a previous study, significant developmental differences in PV expression were observed between primary (V1 and V2), which reached adult levels by 37 days of age, and higher-order visual cortical regions, which reached adult levels by 10 weeks of age in macaques. However, there was no variation in the timing of the plateau among higher-order regions, such as between area TE, 7a, and area 46^18^. The maturation of PV+ cells is recognized to trigger the critical period, a sensitive phase in development^1^. Considering the S-A hierarchical pattern observed in cortical thinning^55,56^ and intracortical myelination^56^, processes that mark the end of critical periods, it is possible that the onset of sensitive periods is widespread across cortex, while their closure follows the S-A hierarchy. However, the finding in the previous study could also be due to the age range, as the model fits could change with the inclusion of younger or older individuals^31,32,57^. For a comprehensive understanding of inhibitory development throughout the lifespan, datasets covering all age ranges from early infancy to adulthood are necessary.

We found that the anesthesia protocol significantly influences the correlation between cell density of inhibitory neurons and the Hurst exponent in mice. Specifically, we observed that the correlation between the Hurst exponent and PV expression vanished completely when medetomidine was used for anesthesia. Typically, resting-state Fmri signals exhibit a frequency distribution with prominent amplitudes at low frequencies (0.01 Hz, 1/f distribution), which is attributed to the slow hemodynamic response to high-frequency neuronal signals. However, vasoconstriction induced by medetomidine anesthesia results in a shorter duration of the hemodynamic response^58^, potentially increasing the likelihood of detecting higher frequency components in the resting-state fMRI signal while diminishing the likelihood of detection of low-frequency components.

Several potential limitations should be noted. First, all datasets included in this study were cross-sectional and of relatively small sample sizes. The mRNA dataset for humans, in particular, only included 1 to 2 children per age and did not include children between the ages of 5 and 7. Additionally, the number of brain regions included was limited, and subcortical regions were not covered. Future work with longitudinal data in a broader age range and including denser sampling of brain regions will be necessary to refine the temporal and spatial sequences of the relationships we report, as well as to better evaluate nonlinearities. Longitudinal data would also make it possible to test relationships between the development of inhibition and functional network specialization. Second, by carefully excluding data with motion artifacts, we may have limited the generalizability of our findings. Most young children move in the scanner, so it is essential to develop more motion-resilient sequences to allow investigators to acquire data from a more representative sample of young kids. Third, the age range of rodent datasets is limited because mice under 30 days old, corresponding to childhood, cannot be scanned due to their small brain size.

In summary, our study builds support for the validity of the Hurst exponent as an inhibitory marker across species. Understanding the developmental processes of inhibition in the human brain is crucial for characterizing both typical and atypical brain development. Inhibitory activity is reduced in autism^28^, schizophrenia^59^, and depression^60^. Additionally, early exposure to substances such as alcohol^61,62^, cocaine^63,64^ can impact inhibitory development. By employing the Hurst exponent as an in vivo marker of inhibition in the developing human brain, we can investigate mechanisms of inhibitory development and guide interventions for risk factors associated with atypical inhibitory development.

## Materials and Methods

### 1. Human data and analyses

#### RNA expression data

Publicly available RNAseq reads per kilobase per million (RPKM) data from the Brainspan Atlas of the Developing Human Brain (https://www.brainspan.org/) were used to characterize patterns of inhibitory marker gene expression across development. RNAseq probes lacking Entrez IDs, which are unique numeric identifiers assigned to genes by the National Center for Biotechnology Information (NCBI) as part of the Entrez Gene database, were excluded (n = 30,984). To address duplicated probes, we retained only the probe with the highest mean expression, resulting in a dataset comprising a total of 21,377 genes, which were subsequently log2 transformed. We used the org.Hs.eg.db package in R to obtain the biological processes regulated by individual genes.

#### fMRI data

The Institutional Review Board at the University of Pennsylvania approved all human fMRI research reported here. Adult data were from a previous study on individual differences in frontoparietal plasticity in humans^65^. Prior to participation, all individuals provided informed, written consent. Participants were recruited through the University of Pennsylvania’s study recruitment system, as well as via community and university advertisements. Inclusion criteria comprised proficiency in English, absence of psychiatric or neurological disorders or learning disabilities, no recent or current illicit substance use, and no contraindications for MRI. A total of 61 participants completed MRI scans. Forty-six adult participants were included in the final sample (M = 21.39 years, SD = 1.91 years; 63% female, 39% male, 0% other or nonbinary; 24% Asian, 33% Black, 17% Hispanic/Latino, 4% Multi-Racial, and 19% White). 77% of participants were undergraduates and 18% were graduate students at the University of Pennsylvania. Exclusions included participants falling asleep during the scans (n = 3), recent illegal substance use not reported during screening but reported during participation (n = 1), inability to tolerate scanning (n = 1), and other technical issues (n = 10).

Child data were collected as part of a study on typical brain development. A subset of these data were previously published (n = 92)^66^. All parents provided informed, written consent for their children’s participation in the study. Verbal assent was obtained from children under eight years old, while those eight years old and older provided written assent. Recruitment efforts targeted Philadelphia and its surrounding areas, using advertisements on public transportation, partnerships with local schools, outreach programs, community family events, and social media ads. Children ranged in age from 4.11 to 10.59 years (M = 6.55, SD = 1.36, 47% male, 53% female, 0% other or nonbinary; 52.9% Black, 42.6% White, 12.5% Asian, and 0.5% other. The final child sample consisted of 128 participants. Exclusions included participants failing to complete the resting-state scan (e.g., due to falling asleep or opting to end the scan prematurely, n = 17) and instances where parents reported a diagnosis of Attention-Deficit/Hyperactivity Disorder or developmental delay during the visit, despite not disclosing it during screening (n = 4). Additionally, motion and quality criteria were applied to mitigate the impact of image quality on our analyses (see fMRI Image Quality and Exclusion Criteria section). These criteria were chosen to balance the need to maximize data inclusion in a young population^67^ with the necessity of minimizing the influence of low-quality data on connectivity metrics^68^.

#### fMRI Image Acquisition

Imaging was performed at the Center for Advanced Magnetic Resonance Imaging and Spectroscopy at the University of Pennsylvania. Scanning was conducted using a Siemens MAGNETOM Prisma 3 T MRI scanner with a Siemens 32-channel coil. Adult participants underwent a T1-weighted multi-echo (MEMPRAGE) structural scan (acquisition parameters: TR = 2530 ms, TI = 1330 ms, TEs = 1.69 ms/3.55 ms/5.41 ms/7.27 ms, BW = 650 Hz/px, 3x GRAPPA, flip angle = 7°, voxel size = 1 mm isotropic, matrix size = 256 × 256 ×176, FOV = 256 mm, total scan time = 4:38) using volumetric navigators^69^ and a five-minute resting-state run (TR = 2000 ms; TEs = 30.20 ms; flip angle = 90°; resolution = 2 mm isotropic) with participants fixating on a cross throughout the scan. Resting-state scanning was continued until at least five minutes of data were acquired with framewise displacement less than 0.5 mm. Children first completed a mock scan to acclimate to typical MRI noises and practice remaining still. During the MRI session, a researcher remained in the scanner room to provide reassurance to the child. Children participated in the same structural and functional scans as adults. Head motion was monitored in real-time using the Framewise Integrated Real-time MRI Monitor system, and 10 minutes of low-motion resting-state data (two runs with framewise displacement < 1 mm) were collected when feasible. For participants with multiple usable resting-state runs, framewise displacement was averaged across runs, weighted by run length. All analyses were adjusted for average framewise displacement and the number of censored volumes.

#### fMRI Image Quality and Exclusion Criteria

The imaging data quality was evaluated through fMRIPrep visual reports and MRIQC 0.14.2 software^70^. Two reviewers manually evaluated all structural and functional images at each preprocessing stage for any image quality concerns. Functional images underwent visual inspection to ensure adequate whole-brain field of view coverage, absence of signal blurring or artifacts, and correct alignment with the anatomical image.

For children’s data, exclusions were made for participants with unusable structural images (n = 1), functional data artifacts (e.g., hair glitter, n = 1), incorrect scanner registration (n = 1), registration failure (n = 18), less than 100 seconds usable data after high-motion outlier volumes (n = 13), and average framewise displacement > 1 mm (n = 6). For participants with multiple usable resting-state runs, framewise displacement was averaged across runs. To ensure that our results were not driven by motion, we conducted an additional analysis with a more stringent motion exclusion criterion. In this pipeline, we excluded participants who had average FD > 0.5 mm (n = 22), for a total of n = 108 participants. None of the adult data met the exclusion criteria of having an average framewise displacement greater than 1 mm.

#### fMRI Image Processing

The T1-weighted (T1w) image underwent intensity nonuniformity correction using N4BiasFieldCorrection^71^ from the Advanced Normalization ToolS (ANTS) version 2.2.0, and used as T1w reference throughout the workflow. Following this, skull stripping was performed using the antsBrainExtraction.sh script (ANTS version 2.2.0), with the OASIS template as the target. The brain mask was refined with a custom variation of the method to reconcile ANTS-derived and FreeSurfer^72^-derived segmentations of the cortical gray matter of Mindboggle (RRID:SCR_002438^73^). Spatial normalization to the ICBM 152 Nonlinear atlases version 2009c. (RRID:SCR_008796^74^) was performed through nonlinear registration with antsRegistration (ANTS, version 2.2.0, RRID:SCR_004757^75^, using brain-extracted versions of both T1w volume and template. Brain tissue segmentation of CSF, white matter (WM), and gray matter was performed on the brain-extracted T1w using fast (Functional MRI of the Brain Software Library (FSL) version 5.0.9; RRID:SCR_002823^76^).

For each resting-state BOLD run, a reference volume and its skull-stripped version were generated using a custom methodology of fMRIPrep version 20.2.0 (RRID:SCR_016216^77^), which is based on Nipype version 1.1.7 (RRID:SCR_002502^78^, nipy/nipype:1.1.7). Brain surfaces were reconstructed using the recon-all command^72^ before other processing, and reconstructed surfaces were used as input to fMRIprep. The BOLD reference was then coregistered to the T1w reference using bbregister (FreeSurfer), employing boundary-based registration^79^. Coregistration was configured with nine degrees of freedom to account for distortions remaining in the BOLD reference. Head-motion parameters with respect to the BOLD reference (transformation matrices and six corresponding rotation and translation parameters) were estimated before any spatiotemporal filtering using the mcflirt tool (FSL version 5.0.9^80^, and slice-time correction was performed using 3dTshift from AFNI 20160207 (RRID:SCR_005927^81^). The BOLD time series were resampled onto the MNIPediatricAsym standard space by applying a single, composite transform, generating a preprocessed BOLD run in MNI152NLin6Asym space.

Several confounding time series were calculated based on the preprocessed BOLD, including FD, DVARS (root-mean-square intensity difference from one volume to the next), and three region-wise global signals (CSF, WM, and the whole brain). FD and DVARS were calculated for each functional run, both using their implementations in Nipype (68). The head-motion estimates calculated in the correction step were also placed within the corresponding confounds file.

All resamplings can be performed with a single interpolation step by composing all the pertinent transformations (i.e., head-motion transform matrices and coregistrations to anatomic and template spaces). Gridded (volumetric) resamplings were performed using antsApplyTransforms (ANTS), configured with Lanczos interpolation to minimize the smoothing effects of other kernels^82^.

Further preprocessing was performed using a confound regression procedure that has been optimized to reduce the influence of participant motion^83,84^; preprocessing was implemented in XCP-D 0.5.0^83^, a multimodal tool kit that deploys processing instruments from frequently used software libraries, including FSL^80^ and AFNI^85^. Further documentation is available at https://xcp-d.readthedocs.io/en/latest/ and https://github.com/PennLINC/xcp_d. Using XCP-D, the preprocessed functional time series on the fsLR cortical surface underwent nuisance regression, using a 36-parameter model that included three global signals (whole brain, CSF, and WM) and six motion parameters, as well as their derivatives, quadratic terms, and derivatives of quadratic terms. Linear regression, implemented in Scikit-Learn (0.24.2), was employed to regress the confounds. Motion censoring was applied, with outlier volumes exceeding FD = 0.3 mm flagged and removed from confound regression. An interpolated version of the BOLD data is created by filling in the high-motion outlier volumes with cubic spline interpolated data, as implemented in nilearn. Any outlier volumes at the beginning or end of the run are replaced with the closest non-outlier volume’s data, in order to avoid extrapolation by the interpolation function. If there was less than 100 seconds of data remaining after removing high-motion outlier volumes, those samples are excluded. No bandpass filtering was performed.

#### Hurst exponent

The estimation of the Hurst exponent uses a discrete wavelet transform and a model of the time series as fractionally integrated processes (FIP) and is estimated using maximum likelihood estimation. FIP model was employed instead of the Fractional Gaussian Noise (fGn) model to eliminate the assumption of stationarity and the upper bound of H = 1. The FIP model provides greater flexibility and potentially increased sensitivity because it offers better estimation of between-subject variability, especially when estimates are close to or surpass H = 1. When H > 1, the time series is deemed non-stationary and displays long memory characteristics, such as being fractal. H is computed using the nonfractal MATLAB toolbox (https://github.com/wonsang/nonfractal)^28,86^. The specific function utilized is bfn_mfin_ml.m function with the ‘filter’ argument set to ‘haar’ and the ‘ub’ and ‘lb’ arguments set to [1.5,10] and [−0.5,0], respectively.

#### Alignment with the S–A axis

We used the S-A axis derived by Sydnor and colleagues^31^. This map encompasses various cortical hierarchies, including functional connectivity gradients, evolutionary cortical expansion patterns, anatomical ratios, allometric scaling, brain metabolism measures, perfusion indices, gene expression patterns, primary modes of brain function, cytoarchitectural similarity gradients, and cortical thickness.

#### GAMs

To flexibly model linear and non-linear relationships among the Hurst exponent and S-A axis rank, we employed Generalized Additive Models (GAMs) using the mgcv package in R^87^. GAMs were employed, with the region-averaged Hurst exponent as the dependent variable, age as a smooth term, and gender, mean framewise displacement and the number of censored as linear covariates. Models were fitted separately for each parcellated cortical region, using thin plate regression splines as the smooth term basis set and the restricted maximum likelihood approach for smoothing parameter selection. The smooth term for age generated a spline representing a region’s developmental trajectory, with a maximum basis complexity (k) set to 3 to prevent overfitting.

For each regional GAM, we assessed the significance of the association between the Hurst exponent and age using an analysis of variance (ANOVA), comparing the full GAM model to a nested model without the age term. A significant result indicates that including a smooth term for age significantly reduced the residual deviance, as determined by the chi-squared test statistic. For each regional GAM, we identified the specific age range(s) where the Hurst exponent significantly changed using the gratia package in R. Age windows of significant change were determined by examining the first derivative of the age smooth function (Δ Hurst exponent/Δ age) and assessing when the simultaneous 95% confidence interval of this derivative did not include 0 (two-sided). To quantify the overall magnitude and direction of the association between the Hurst exponent and age, referred to as a region’s overall age effect, we calculated the partial R^2^ between the full GAM model and the reduced model for effect magnitude. We then signed the partial R^2^ based on the average first derivative of the smooth function for effect direction. We sorted 400 parcels based on their S-A rank and divided them into three groups: sensory, middle and association regions. Subsequently, we performed the same GAM analysis for the averaged values across 133 parcels within each group.

#### Spin-based spatial permutation testing

To address distance-dependent spatial autocorrelation in cortical data, we evaluated the significance of Pearson’s correlations between two whole-brain cortical feature maps (the Hurst exponent) using non-parametric spin tests. These tests, also known as spatially constrained rotation tests, compare the observed correlation to a null distribution obtained by spatially iterating (spinning) one of the feature maps. Specifically, the null distribution is generated by rotating spherical projections of one map while preserving its spatial covariance structure. The P value (P_spin_) is determined by comparing the empirical correlation to the distribution obtained from 10,000 spatial rotations. Spin tests were conducted using the spin permutation test algorithm available in the ENIGMA toolbox for Python (https://enigma-toolbox.readthedocs.io/en/latest/pages/08.spintest/index.html).

### 2. Mouse data and analyses

#### Inhibitory neuron cell density

For cell density analysis of PV+, SST+, and VIP+ cells, we used the dataset published in Kim et al., 2017^5^. Five male and five female transgenic mice were used per each cell type. Cell counting and distribution mapping were conducted using a platform developed by Kim et al., 2015^5^, which relies on automated imaging via serial two-photon tomography (STPT) and data analysis through machine learning algorithms. Within this process, automatic cell counting within specific anatomical regions was achieved by tallying the number of registered signals within each anatomical label using custom-built codes. Cell counting was carried out in evenly spaced and partially overlapping 3D sphere voxels (with a diameter of 100 μm and 20 μm apart) to accurately digitize and visualize the distribution of detected signals throughout the entire brain in an unbiased manner.

#### fMRI data

For fMRI analysis, a publicly available fMRI dataset (doi:10.18112/openneuro.ds001653.v1.0.2) was used. All applicable international, national, and/or institutional guidelines for the care and use of animals were followed. All procedures performed in studies involving animals were in accordance with the ethical standards of the Institutional Animal Care and Use Committee (A*STAR Biological Resource Centre, Singapore, IACUC #161134/171203). For the analysis of correlation between cell density and the Hurst exponent, 10 Female C57BL/6 mice (Janvier, Le Genest-St Isle, France), aged three months were included. A total of 57 runs were conducted, with each mouse undergoing six runs over two sessions. The mice were housed in standard mouse cages under a 12-hour light/dark cycle, with access to food and water ad libitum. Anesthesia was induced with 4% isoflurane. Subsequently, the animals were placed on an MRI-compatible cradle while isoflurane was reduced and kept to 1%. Care was taken to maintain the animal temperature at 37°C. Another 10 C57BL/6 female mice (Janvier, Le Genest-St Isle, France), aged three months were included for testing the correlation under Med/Iso (Medetomidine 0.05 mg/kg bolus/0.1 mg/kg/h infusion (i.v.) + Isoflurane 0.5%) anesthesia protocol. A total of 57 runs were conducted, with each mouse undergoing six runs over two sessions. For those mice, anesthesia was induced with 4% isoflurane. Subsequently, a bolus with a mixture of medetomidine (Dormitor, Elanco, Greenfield, Indiana, USA) and Pancuronium Bromide (muscle relaxant, Sigma-Aldrich Pte Ltd, Singapore) was administered subcutaneously (0.05 mg/kg), followed by a maintenance infusion (0.1 mg/kg/h) while isoflurane was simultaneously reduced and kept to 0.5%.

To analyze the developmental trajectory of the Hurst exponent, a total of 72 wild-type (comprising 40 females and 32 males) within the age range of one to eight months were included, distributed across four distinct age groups: 1 month (n = 21, with 9 females and 12 males), 2 months (n = 16, with 6 females and 10 males), 5 months (n = 17, with 13 females and 4 males) and 8 months (n = 18, with 12 females and 6 males). The mice were housed in standard mouse cages under a 12-hour light/dark cycle, with access to food and water ad libitum. Anesthesia was induced using isoflurane (Abbott) in a 1:4 oxygen to air mixture, with a concentration of 3% for induction and 1.4% during measurements, administered via a facemask. During the experiments, the mice were positioned on a mouse support apparatus with ear bars used to minimize motion. Body temperature was continuously monitored using a rectal thermometer and maintained at 36.5°C ± 0.5°C through an adjustable warm water bath integrated into the support. The entire procedure, including mouse preparation, typically lasted for 45 minutes.

#### fMRI Image Acquisition

Data were acquired on an 11.75 T scanner (Bruker BioSpin MRI, Ettlingen, Germany) equipped with a BGA-S gradient system, a 72 mm linear volume resonator coil for transmission and a cryoprobe 2 × 2 phased-array surface coil. Images were acquired using Paravision 6.0.1 software. An anatomical reference scan was acquired using a spin-echo Turbo-RARE sequence: field of view (FOV) = 17 × 9 mm^2^, FOV saturation slice masking non-brain regions, number of slices = 28, slice thickness = 0.35, slice gap = 0.05 mm, matrix dimension (MD) = 200 × 100, repetition time (TR) = 2742 ms, echo time (TE) = 30 ms, RARE factor = 8, number of averages = 2. fMRI was acquired using a gradient-echo echo-planar imaging (GE-EPI) sequence with the same geometry as the anatomical scan: MD = 90 × 60, in-plane resolution = 0.19 × 0.15 mm2, TR = 1000 ms, TE = 11.7 ms, flip angle = 50°, volumes = 180, bandwidth = 119047 Hz. Field inhomogeneity was corrected using the MAPSHIM protocol.

#### fMRI Image Processing

A study-specific template was created from the individual anatomical images using the buildtemplateparallel.sh script from the Advanced Normalization Tools (ANTs, 20150828 builds^88^). The study template was subsequently registered to the reference template of the Allen Mouse Common Coordinate Framework version 3 (CCF v3)^89–91^. Individual anatomical images were corrected for b1 field inhomogeneity (N4BiasFieldCorrection), denoised (DenoiseImage), intensity thresholded (ImageMath), brain masked (antsBrainExtraction.sh), and registered to the study template, resampled to a 0.2 × 0.2 × 0.2 mm3 resolution, using a SyN diffeomorphic image registration protocol (antsRegistration). The functional EPI time series were despiked (3dDespike) and motion-corrected (3dvolreg) using the Analysis of Functional NeuroImages software (AFNI_16.1.15^85^). A temporal average was estimated using fslmaths (FMRIB Software Library 5.0.9^92^), corrected for the b1 field (N4BiasFieldCorrection), brain masked (antsBrainExtraction.sh), and registered to the anatomical image (antsRegistration). The fMRI time series were bandpass filtered (3dBandpass) to a 0.01 to 0.25 Hz band. An independent component analysis (MELODIC) was performed and used as a basis to train a FIX classifier^93^ in 15 randomly selected runs. The classifier was applied to identify nuisance-associated components that were regressed from the fMRI time series. The EPI to anatomical, anatomical to study template, and study template to Allen Mouse CCF v3 registrations were combined into one set of forward and backward transforms (ComposeMultiTransform). The combined transforms were applied to the denoised fMRI data. Signal time series for each region of interest (ROI) in the Allen Mouse Brain atlas^89^ were extracted.

#### Hurst exponent

We used the same metrics as those applied to humans to calculate the Hurst exponent for mice.

### 3. Rat data and analysis

#### fMRI data

For the fMRI analysis, we utilized the Standard Rat dataset as described in Grandjean et al., 2023(40). All fMRI acquisitions were conducted following approval from the relevant local and national ethics authorities. To investigate the developmental trajectory of the Hurst exponent, we included a total of 145 rats, comprising 54 females and 91 males, aged between one and four months. These rats were distributed across four distinct age groups: 1.0 month (n = 10, males only), 1.5 months (n = 10, males only), 2.0 months (n = 68, with 26 females and 42 males), 2.5 months (n = 20, with 5 females and 15 males), 2.9 months (n = 3, females only), 3.0 months (n = 14, with 11 females and 3 males), and 4.0 months (n = 20, with 9 females and 11 males).

#### fMRI Image Acquisition

Rats were anesthetized using 4% isoflurane and a subcutaneous (s.c.) bolus of 0.05 mg/kg medetomidine for induction, followed by maintenance anesthesia with 0.4% isoflurane and a continuous infusion of 0.1 mg/kg/h medetomidine s.c. Imaging was conducted using a gradient-echo echo-planar imaging technique 40 minutes after anesthesia induction. The imaging parameters included a repetition time of 1,000 ms, echo time/flip angle/bandwidth defined according to field strength, 1,000 repetitions, a matrix size of 64 × 64, a field of view of 25.6 × 25.6 mm^2^, and 18 interleaved axial slices of 1 mm thickness with a 0.1-mm gap. The full protocol can be found at https://github.com/grandjeanlab/StandardRat.

#### fMRI Image Processing

The scans were organized following the BIDS format. Pre-processing was conducted separately for each scan session using a reproducible containerized software environment for RABIES 0.3.5 (Singularity 3.7.3–1.el7, Sylabs). This involved several steps: autobox, N4 inhomogeneity correction, motion correction, rigid registration between functional and anatomical scans, non-linear registration between anatomical scan and template, and resampling to a common space at 0.3 × 0.3 × 0.3 mm^3^ resolution. To address differences in brain size, image contrast, and susceptibility distortions in rodent images, a workflow for volumetric image registration and brain masking was developed. Quality control was ensured through visual inspection of pre-processing outputs for all scans. Global signal regression were performed along with motion regression, spatial smoothing to 0.5 mm^3^, a high-pass filter at 0.01 Hz, and a low-pass filter at either 0.1 Hz or 0.2 Hz. The ICA-AROMA method was adapted from humans to rats by utilizing dedicated rat cerebrospinal fluid and brain edge masks and by training the classifier parameters based on a set of rodent images. Visual inspection of the components and their classifications showed that less than 5% of the plausible signal components were mistakenly labeled as noise. Signal time series for each region of interest (ROI) in the SIGMA anatomical atlas(42) were extracted using the NiftiLabelsMasker function in Nilearn 0.7.1.

#### Hurst exponent

We used the same metrics as those applied to humans to calculate the Hurst exponent for rats.

## Author Contributions and Notes

M.N., M.K., and M.M. conceptualized the study; M.N., E.Y., and M.K. M.N. and A.P.M. conceptualized the study; C.L.M, A.T.P., U.A.T., A.L.B. collected data; M.N. analyzed the data and prepared the figures; M.N., M.E, M.W. and A.P.M. interpreted the results; M.N. and A.P.M. wrote the manuscript; A.P.M. and J.G. supervised M.N; all authors provided feedback on the manuscript. The authors declare no conflict of interest.

## Acknowledgment

First, we thank all of the individuals who participated in this research. We also thank Maayan Ziv, Steven Lopez, Cassandra Garcia, Priya Deliwala, Jasmine Forde, Katrina Simon, Sophie Sharp, Yoojin Hahn, Stephanie Bugden, Jamie Bogert, Alexis Broussard, Ava Cruz, Samantha Ferleger, Destiny Frazier, Jessica George, Abigail Katz, Sun Min Kim, Hunter Liu, Dominique Martinez, Ortal Nakash, Emily Orengo, Christina Recto, Leah Sorcher, Alexis Uria, Adam Pines, Quentin Wedderburn, Connor Kendzora, Anna Meaney, Leah Sorcher, Aparna Ramanujam, Daniel Southwick, Alexis Broussard and Andrea Gomez for their help with data acquisition and cleaning. This study was supported by a National Science Foundation CAREER award (to A.P.M. under Grant No. 2045095), National Institute on Drug Abuse Grant 1R34DA050297-01 (A.P.M.), Jacobs Foundation Early Career Award (A.P.M.), CIFAR Global Azrieli Fellowship (to A.P.M), Quad Fellowship (to M.N.), Nakajima Foundation Scholarship (to M.N.), National Science Foundation Graduate Research Fellowships (to A.L.B., U.A.T. and C.L.M. under Grant No. DGE-1845298), Behavioral and Cognitive Neuroscience Training Grant (to A.T.P. under Grant No. NIH T32-MH017168), as well as start-up funds from the University of Pennsylvania.

