## Supplementary Material for "The Hurst exponent as a marker of inhibition in the developing brain"

### Supporting Information for
The Hurst exponent as a marker of inhibition in the developing brain

Monami Nishio[1], Monica E. Ellwood-Lowe[1], Mackenzie Woodburn[1], Cassidy L. McDermott[1], Anne T. Park[1], Ursula A. Tooley[1], Austin L. Boroshok[1], Joanes Grandjean[2,3], Allyson P. Mackey[1]*

Allyson Mackey


**This PDF file includes:**

Figures S1 to S5

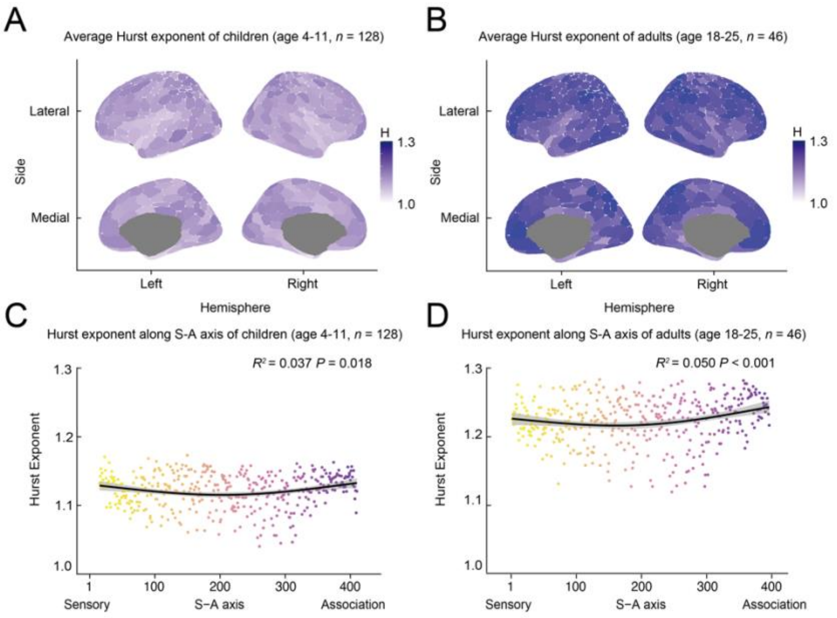


**Supplementary Figure 1. The Hurst exponent across cortex.** We parcellated the brain regions with Schaefer's set of 400 delineations[33]. In order to characterize how the Hurst exponent varies along the sensory-association (S-A) axis[31], we took the Hurst exponent from 400 parcels, arranged based on the S-A axis. **(A, B)** Group-averaged Hurst exponent maps for children (ages 4-11 years, *n* = 128) (A) and adults (ages 18-25 years, *n* = 46) (B). **(C, D)** Mapping of the Hurst exponent along the S-A axis for children (C) and adults (D). The GAM-predicted Hurst exponent value at each parcel along the S-A axis is displayed with a 95% confidence interval ($R^2$ = 0.037, $P$ = 0.018 for children; $R^2$ = 0.050, $P$ < 0.001 for adults). As expected, adults exhibited significantly higher Hurst exponents compared to children throughout the cortex, suggesting elevated levels of inhibition in adults relative to children. (children vs adults, independent t-test, $t(172)$ = 4.301, $P$ < 0.001). However, it is likely that this observed difference can be partially attributed to variations in head motion. After controlling for age and in-scanner head motion, the gender difference did not survive FDR correction for any of the parcels in children or adults.

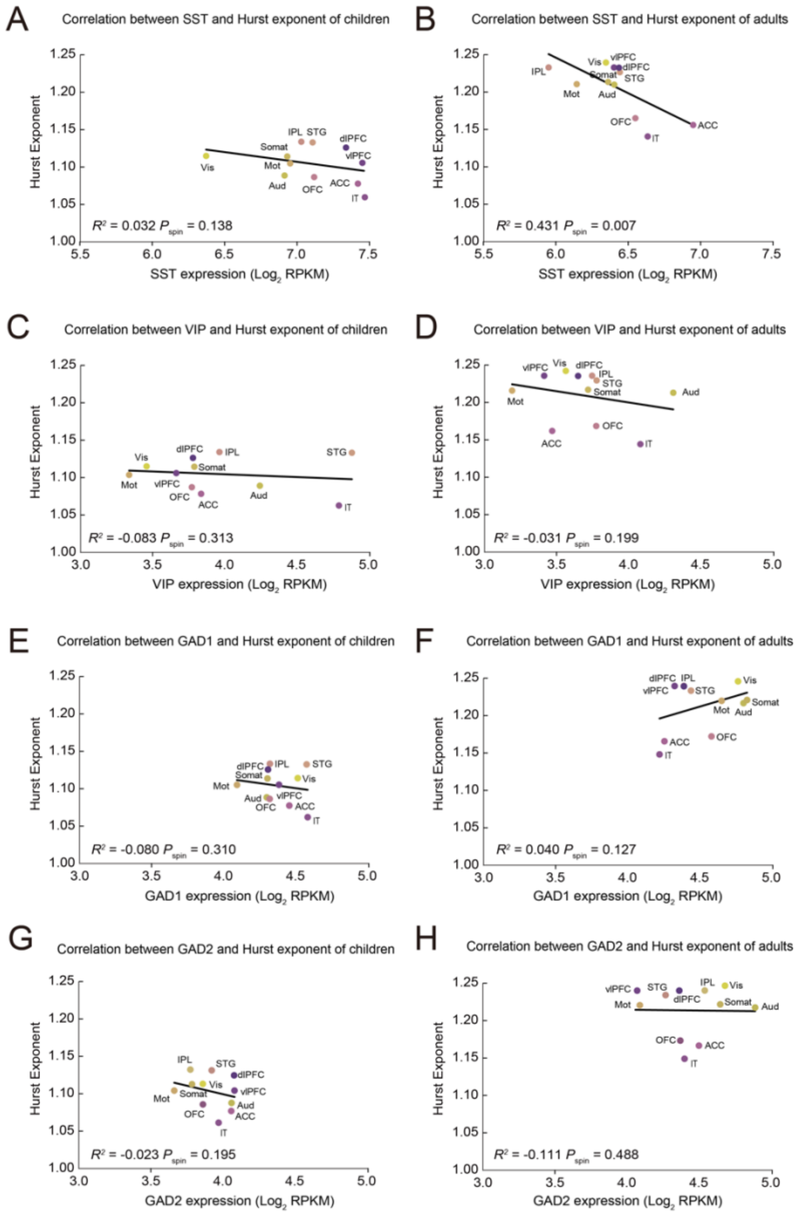


**Supplementary Figure 2. Correlations between cell density and the Hurst exponent in humans.** **(A, B)** Correlation between SST expression and the Hurst exponent in children (A, $R^2$ = 0.032, $F(1, 9)$ = 1.331, $P_{spin}$ = 0.138) and adults (B, $R^2$ = 0.431, $F(1, 9)$ = 8.570, $P_{spin}$ = 0.007). **(C, D)** Correlation between VIP expression and the Hurst exponent in children (C, $R^2$ = -0.083, $F(1, 9)$ = 0.236, $P_{spin}$ = 0.313) and adults (D, $R^2$ = -0.031, $F(1, 9)$ = 0.701, $P_{spin}$ = 0.199). **(E, F)** Correlation between GAD1 expression and the Hurst exponent in children (E, $R^2$ = -0.080, $F(1, 9)$ = 0.258, $P_{spin}$ = 0.310) and adults (F, $R^2$ = 0.040, $F(1, 9)$ = 1.412, $P_{spin}$ = 0.127). **(G, H)** Correlation between GAD2 expression and the Hurst exponent in children (G, $R^2$ = -0.023, $F(1, 9)$ = 0.772, $P_{spin}$ = 0.195) and adults (H, $R^2$ = -0.111, $F(1, 9)$ = 0.002, $P_{spin}$ = 0.488).

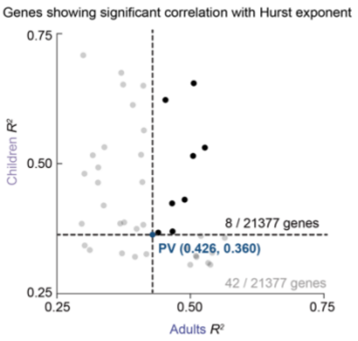


**Supplementary Figure 3. mRNA markers showing a significant correlation with the Hurst exponent. (A)** The x-axis shows the $R^2$ value for the correlation between mRNA expression in adults (ages 18-23 years, $n$ = 4) and the Hurst exponent in adults (ages 18-25 years, $n$ = 46). The y-axis shows the $R^2$ value for the correlation between mRNA expression in children (ages 3-13 years, $n$ = 6) and the Hurst exponent in children (ages 4-11 years, $n$ = 128). There were 42 out of 21377 genes (gray dots) that exhibited a significantly positive correlation with both adults and children's Hurst exponents. Among them, eight genes (black dots) have a higher $R^2$ than PV for both adults and children.

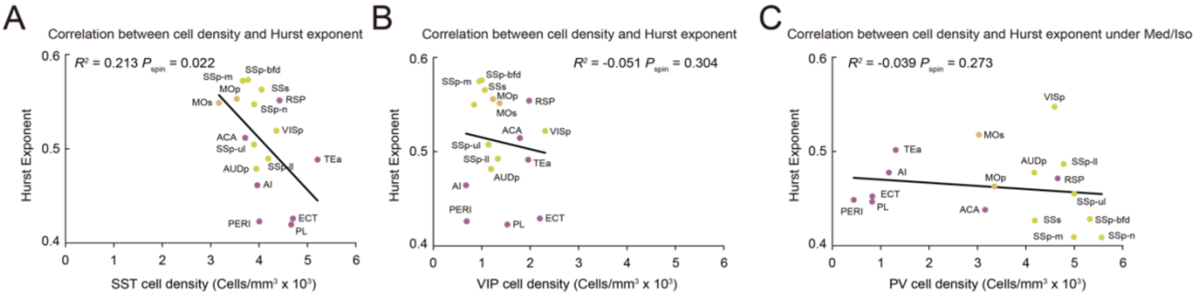


**Supplementary Figure 4. The correlation between inhibitory cell densities and the Hurst exponent in mice. (A)** Correlation between the Hurst exponent and the density of SST+ cells. Linear regression analysis indicates a significant correlation ($R^2$ = 0.213, *F(1, 15) = 5.326*). (B) Correlation between the Hurst exponent and the density of VIP+ cells ($P_{spin}$ = 0.022, B; $R^2$ = -0.051, *F(1, 15)* = 0.229,$P_{spin}$ = 0.304). **(C)** Correlation between PV+ cell density and the Hurst exponent under Med/Iso anesthesia protocol. The linear regression analysis shows a non-significant correlation ($R^2$ = -0.039, *F(1, 15)* = 0.400, $P_{spin}$ = 0.273).

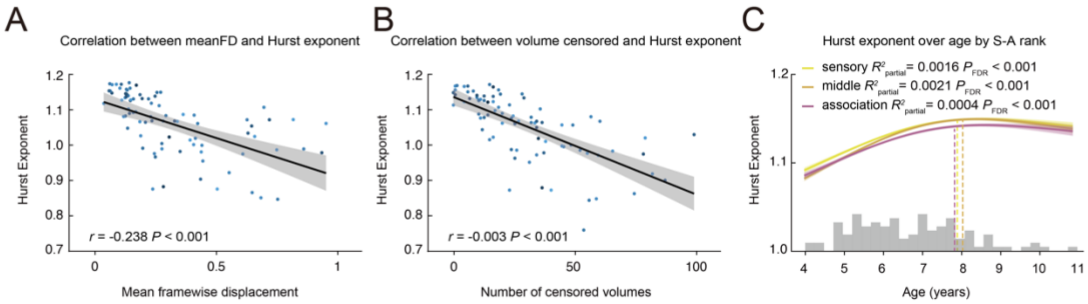


**Supplementary Figure 5. Sensitivity analysis. (A)** Correlation between mean framewise displacement and the Hurst exponent across samples in children while controlling for age ($r = -0.238$, $F(2, 125) = 16.310$, $P < 0.001$). The negative linear fit between these measures is shown with a 95% confidence interval. **(B)** Correlation between number of censored volumes and the Hurst exponent across samples in children while controlling for age ($r = -0.003$, $F(2, 125) = 36.730$, $P < 0.001$). The negative linear fit between these measures is shown with a 95% confidence interval. **(C)** GAM-predicted developmental trajectories of the Hurst exponent. The 400 parcels are segmented along the S-A axis into three bins, each containing 133 parcels. Regional trajectories for each bin illustrate the GAM-predicted Hurst exponent values at each age, accompanied by a 95% error band based on standard errors. A dashed line marks the age at which the first derivative of the age smooth function (Δ Hurst exponent / Δ age) becomes insignificant. Sensory; partial $R^2 = 0.0016$, $P_{FDR} < 0.001$, middle; partial $R^2 = 0.0021$, $P_{FDR} < 0.001$, association; partial $R^2 = 0.0004$, $P_{FDR} < 0.001$.
